# Spatial organisation and homeostasis of epithelial lineages at the gastroesophageal junction is regulated by the divergent Wnt mucosal microenvironment

**DOI:** 10.1101/2021.08.05.455222

**Authors:** Naveen Kumar, Rajendra Kumar Gurumurthy, Pon Ganish Prakash, Shilpa Mary Kurian, Christian Wentland, Volker Brinkmann, Hans-Joachim Mollenkopf, Tobias Krammer, Christophe Toussaint, Antoine-Emmanuel Saliba, Matthias Biebl, Christian Juergensen, Bertram Wiedenmann, Thomas F Meyer, Cindrilla Chumduri

**Affiliations:** Department of Microbiology, University of Würzburg, Würzburg, Germany; Department of Molecular Biology, Max Planck Institute for Infection Biology, Berlin, Germany; Surgical Clinic Campus Charité Mitte, Charité University Medicine, Berlin, Germany; Department of Hepatology and Gastroenterology, Charité University Medicine, Berlin, Germany; Institute for RNA-based Infection Research (HIRI), Helmholtz Center for Infection Research (HZI), Würzburg, Germany; Laboratory of Infection Oncology, Institute of Clinical Molecular Biology (IKMB), Christian Albrechts University of Kiel, Kiel, Germany

## Abstract

The gastroesophageal junction (GEJ), where squamous and columnar epithelia meet, is a hotspot for Barrett’s metaplasia development, dysbiosis and carcinogenesis. However, the mechanisms regulating GEJ homeostasis remain unclear. Here, by employing organoids, bulk and single-cell transcriptomics, single-molecule RNA in situ hybridisations and lineage tracing, we identified the spatial organisation of the epithelial, stromal compartment and the regulators that maintain the normal GEJ homeostasis. During development, common KRT8 progenitors generate committed unilineage p63/KRT5-squamous and KRT8-columnar stem cells responsible for the regeneration of postnatal esophagus and gastric epithelium that meet at GEJ. A unique spatial distribution of Wnt regulators in the underlying stromal compartment of these stem cells creates diverging Wnt microenvironments at GEJ and supports their differential regeneration. Further, we show that these tissue-resident stem cells do not possess the plasticity to transdifferentiate to the other lineage with the altered Wnt signals. Our study provides invaluable insights into the fundamental process of GEJ homeostasis and is crucial for understanding disease development.

## Introduction

The mucosal epithelium of different organs often exposed to extrinsic factors like diet, toxins, and microbes are predisposed to carcinogenesis. Epithelial transition zones where two different epithelial types meet represent hotspots of preneoplastic metaplasia, altered microbiota and cancer development ^1–4^. Gastroesophageal junction (GEJ) is one of those transition zones defined by the Z-line where the mucosa of the distal esophagus and the proximal stomach meet. This anatomical structure acts as a sphincter and is critical for barrier function, including preventing stomach contents from refluxing upward into the esophagus. Failure in the anti-reflux function of the GEJ leads to gastroesophageal reflux disease (GERD), a condition in which acidic stomach contents moves into the esophagus, damaging the esophageal mucosa ^5^. GERD patients often develop Barrett’s metaplasia (BE), a precursor of esophageal adenocarcinomas, characterised by the replacement of stratified squamous epithelium with glandular or intestinal-type epithelium in the esophagus. Further increasing obesity and altered microbiota trends are implicated as additional risk factors of BE and carcinogenesis ^6–8^. Due to the deadly nature and fast-increasing incidence of GEJ adenocarcinomas accounting for a 6-fold increase during the past four decades, with a five-year survival of 15%, have gained the attention of clinicians and researchers ^9–12^.

Recently, several studies presented different hypotheses for the cell of origin of BE at GEJ. These include mechanisms of transdifferentiation of the squamous esophageal epithelium to BE ^13–15^, circulating bone marrow stem cells ^16^, unique KRT7+ residual embryonic stem cells ^17^, or transitional basal epithelial cells p63+/KRT5+/KRT7+ ^18^ between esophagus and stomach epithelium at the GEJ, LGR5 cells from cardia region or the first gland of the stomach ^1,19^ and submucosal glands of the esophagus ^20,21^. However, despite these studies, epithelial lineages and mechanisms involved in epithelial regeneration, the establishment of the squamocolumnar epithelial boundary at normal adult GEJ and the role of stromal microenvironment in their homeostasis remian unknown. Clearly, Wnt signalling is essential for regulating the gastrointestinal tract homeostasis, stem cell proliferation and differentiation ^22,23^. In addition, Wnt pathways are implicated in cancer development in the stomach and esophagus ^24–26^, and dysregulation of the Wnt pathway is associated with BE development ^27^.

This study unravelled the epithelial cell types, their spatial organisation and plasticity and identified the Wnt mucosal microenvironment as a critical regulator of the squamocolumnar epithelial border homeostasis at the GEJ. During embryogenesis, the bipotent KRT8+ primitive epithelium lining of the gut mucosa differentiates into postnatal unilineage squamous and columnar epithelial stem cells. By employing mice and patient-derived organoids, lineage tracing, immunostaining and single-molecule in situ RNA hybridisation (smRNA-ISH), we found that these distinct lineage-specific stem cells are responsible for the regeneration of squamous and columnar epithelia at the GEJ. Unique spatial distribution of the Wnt regulators from the stroma underlying squamous and columnar epithelium creates a distinct Wnt inhibitory or activating microenvironment driving their regeneration and thus establishing the epithelial boundary at GEJ. Lineage tracing confirmed that Wnt signalling is critical for the differential proliferation of these distinct lineage-specific stem cells but does not drive transdifferentiation to other lineages. Bulk and single-cell sequencing of squamous and columnar organoids revealed epithelial subpopulations and molecular signatures recapitulating the in vivo squamous stratified esophageal and columnar stomach epithelium and their functions. Thus, a diverging Wnt microenvironment at GEJ establishes the borders between distinct epithelial stem cell lineages that possess different physiological functions; however, it is not involved in transdifferentiation into columnar or intestinal-type BE. These insights have implications in understanding the largely unknown mechanisms of tissue response to damage during repair and the mechanisms that contribute to metaplasia and cancer development.

## Results

### Epithelial cell types and tissue microenvironment at the gastroesophageal junction

The adult human esophageal mucosa is lined with stratified squamous epithelium that meets the glandular columnar epithelium lined stomach at the gastroesophageal junction (GEJ) (Figure 1A). Whereas in the mouse, the esophagus opens into the stomach that comprises two regions- a glandular stomach and stratified squamous epithelium lined fore-stomach similar to the esophagus (Figure 1A). To gain insights into the epithelial stem cells involved in establishing the adult GEJ, we analysed the mucosal lining of the GEJ at different embryonic and adult stages. Tissue sections were made through the esophagus and entire stomach mucosa from embryonic day 13 (E-13), E-16 and E-19 and fluorescence immunohistochemistry were performed for the transcription factor p63, a regulator of stratified squamous epithelium and cytokeratins KRT5, KRT8, and KRT7 (Figure 1B, Supplementary Figure 1A). On E-13, the entire stomach region consisting of simple columnar epithelial cells were labelled with KRT7 and KRT8. Further, multilayered squamous epithelial cells expressing p63 without KRT5 expression appeared from the proximal esophagus to the distal region, below the KRT7+/KRT8+ simple columnar epithelium (Figure 1B-i, Supplementary Figure 1A-i). On E-16, the entire mucosa of the esophagus and forestomach was lined by p63+ squamous epithelial cells with a faint expression of KRT5 below the KRT7+KRT8+ columnar cells. Notably, near the junction, KRT7+KRT8+ precursor cells show differentiation into KRT7+KRT8+P63+KRT5- and these cells position as subcolumnar cells (Figure 1B-ii, Supplementary Figure 1A-ii). By E-19, these KRT7+KRT8+P63+KRT5-cells subsequently gain KRT5 expression but lose KRT8/KRT7 expression. The KRT7+KRT8+ expressing cells above the P63+KRT5+ cells slough off, thus visibly demarcating the squamous and glandular regions of the esophagus and stomach (Figure 1B-iii, Supplementary Figure 1A-iii). In the adult mouse, the squamous cells in the esophagus were KRT5+P63+/KRT8-KRT7- and columnar cells in the stomach were KRT5−P63−/KRT8+KRT7+ (Figure 1C, D). Similar cytokeratin patterns were confirmed in the human esophagus and stomach epithelium meeting at GEJ (Figure 1H, Supplementary Figure 1B). Furthermore, the smRNA-ISH analysis revealed that *Krt5* mRNA is specifically expressed in the esophageal epithelia but not in the columnar epithelium of the stomach (Figure 1E), while *Krt8* mRNA is highly expressed exclusively in the columnar epithelium of the stomach (Figure 1F). In contrast to the previous reports that a few unique KRT7+ embryonic progenitor cells are retained at adult GEJ and are the precursors of BE ^17,18^, we observed that *Krt7* expression is not confined only to the junctional region. Instead, the *Krt7* gene is highly expressed in the entire columnar stomach epithelium and, to a lesser extent, also in the basal cells of the esophagus (Figure 1G). Thus, postnatal GEJ comprises two major cell types, squamous stratified KRT5+P63+ epithelial cells of the esophagus joining the KRT7+KRT8+ columnar cells of stomach cells.

**Figure 1.**
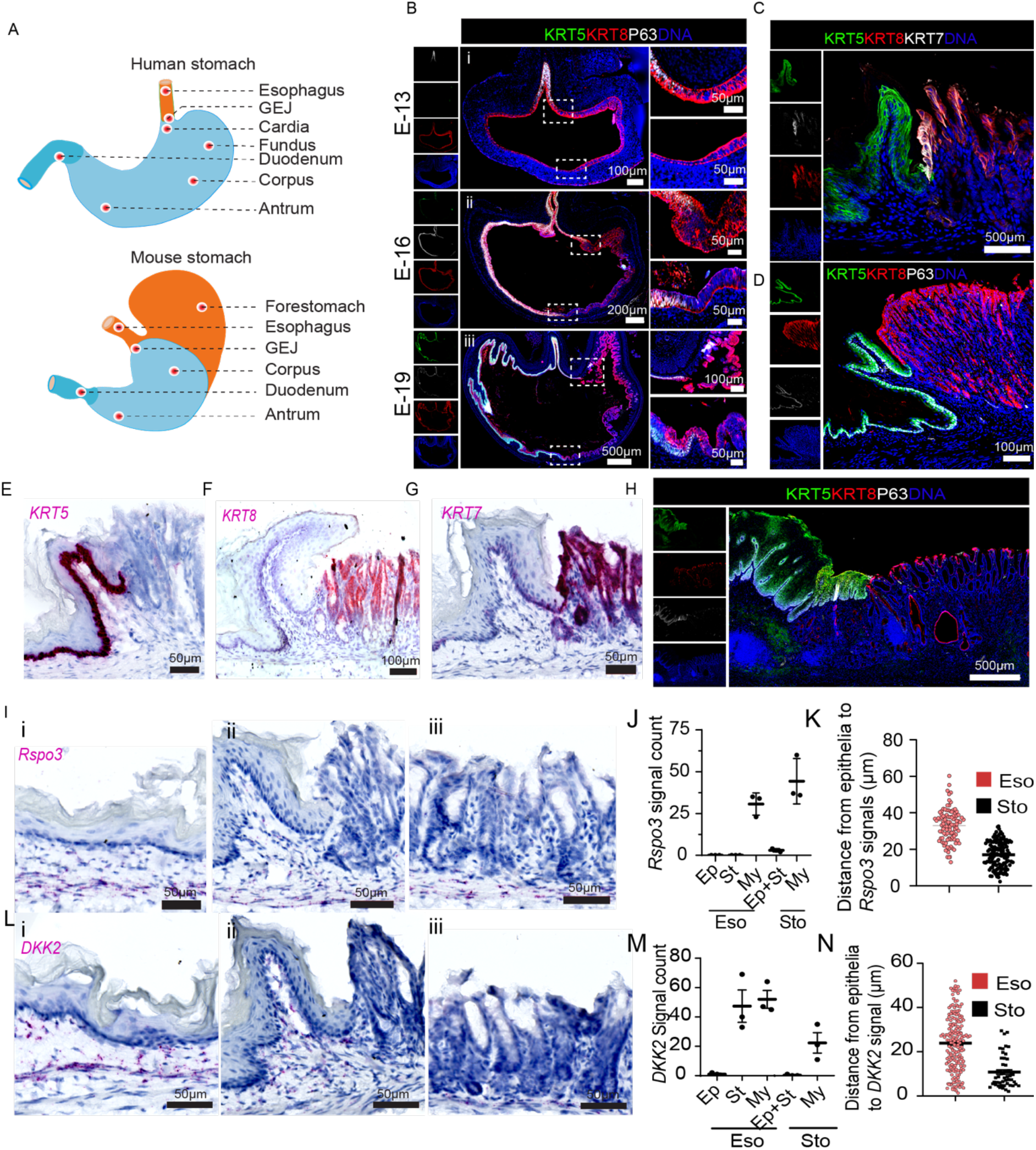
Epithelial cell types and stromal microenvironment define the gastroesophageal junction’s spatial organisation. (A) Schematic of human and mouse esophagus and stomach anatomy, including gastroesophageal junction (GEJ). GEJ in postnatal humans is formed where the distal squamous stratified epithelium lined esophagus joins the proximal columnar epithelium of the stomach (cardia). In mouse stratified epithelium lines, the esophageal and forestomach that opens into columnar epithelium lined stomach forming GEJ. (B) Tiled images of tissue sections from stomachs of embryonic day 13 (E-13) (i), E-16 (ii) and E-19 (iii) mice stained with KRT5 (green), KRT8 (Red), P63 (white) and nuclei stained with DAPI (blue). A magnified view of the boxed GEJ regions is shown in the right panel. (C-D) Tiled images of adult mouse GEJ tissue stained with KRT5 (green), KRT8 (Red), KRT7 (white) (C) and KRT5 (green), KRT8 (Red), P63 (white) (D) nuclei were stained with DAPI (blue). (E-G) smRNA-ISH images of mouse GEJ tissue probed with *Krt5*, *Krt8* and *Krt7*. Nuclei are labelled with blue. (H) Tiled image of adult human GEJ tissue sections stained with KRT5 (green), KRT8 (Red), P63 (white), and nuclei stained with DAPI (blue). (I-N) smRNA-ISH images for the Wnt pathway genes *Rspo3* (I) and *Dkk2* (L) in the mouse esophagus tissue (i), GEJ (ii), and stomach glands (iii). Nuclei are labelled with blue. Quantification of *Rspo3* (J) and *Dkk2* (M) signal counts in Epithelia (Ep), stroma (St), and Myofibroblast (My) in the mouse GEJ tissue regions, distance (μm) from epithelia to *Rspo3* (K) and *Dkk2* signal (N). Signal counts were performed from three non-overlapping 100 μm^2^ areas of the image. Images are representative of n=3 mice or human donors. Tiled images shown in B-I, L were acquired with an AxioScan imager.

Glandular epithelium of the stomach and its regeneration is regulated by extrinsic and cell-autonomous Wnt signalling ^23,28,29^. However, the role of Wnt signalling in the esophagus and at GEJ is not known. Thus, we performed spatial expression analysis of genes that function as agonistic and antagonistic morphogens of the Wnt pathway in the mouse GEJ tissue. R-spondin-3 (*Rspo3)*, which potentiates the Wnt signalling expressed in the myofibroblast (Myo) in both the esophagus and stomach tissue (Figure 1I-J). However, the proximity of *Rspo3* signals to the stem cell compartment of the esophagus and stomach differed. In the stomach, myofibroblasts are located proximal to the stem cells of the gastric glands, while in the esophagus, the stromal region separates basal epithelial stem and myofibroblast cells. Thus, the average distance of the *Rspo3* signal to the epithelia is greater in the esophagus than in the stomach (Figure 1J-K). Further, Wnt pathway inhibitor *DKK2* ^30^ is highly expressed in the stroma and myofibroblast cells of the esophagus and to a significantly lesser extent in myofibroblast cells in the stomach (Figure 1L-N). Thus, the squamous and columnar epithelium at GEJ are associated with spatially defined distinct Wnt microenvironments.

### Organoids of gastric and esophageal epithelium reveal distinct Wnt dependency

Based on the above-observed distribution of Wnt signals in the microenvironment (Figure 1I-N), we tested the role of Wnt signalling in stemness and regeneration of gastric and esophageal epithelial types by employing organoid technology. We isolated the primary cells from the mouse esophagus and stomach, and organoids were grown in the presence and absence of WNT3a, RSPO1 (W/R) containing media in addition to mEGF, mNoggin, FGF10, Nicotinamide, Forskolin, and Alk3/4/5 inhibitor A83-01. Mouse esophageal stem cells grew into mature squamous stratified esophageal epithelial organoids in both presence and absence of WNT3a and RSPO1 (Figure 2A), suggesting that Wnt signalling is non-essential for the esophageal organoids formation. In contrast, as previously described ^29,31^, Wnt signalling is essential for stomach organoid growth as the presence of WNT3a and RSPO1 conditioned media was necessary for the formation of stomach organoids (Figure 2A). Strikingly, in the case of humans, the presence of WNT3a and RSPO1 showed an inhibitory effect on esophageal organoid growth, while their absence supports the growth (Figure 2E). Esophageal organoids contain multi-layered epithelium as opposed to stomach organoids which consist of the single-layered columnar epithelium with a hollow centre (Figure 2B, F). To determine the long-term growth efficiency of esophageal organoids in the presence and absence of (W/R) media, the percentage of organoid formation was quantified at passages 8 and 12. In the presence of WNT factors, esophageal organoid formation efficiency decreased from passage 8 to 12, as opposed to WNT deficient media (Figure 2C). Esophageal organoids were able to grow more than 22 passages in the absence of (W/R) media, whereas these organoids ceased to grow at passage 13 when cultured in the presence of WNT factors (Figure 2D). Thus, the Wnt signalling factors are not essential for the establishment, long-term culturing and expansion of esophageal squamous epithelial organoids as opposed to stomach columnar organoids. Cultured organoids maintained epithelial lineage specificity and morphology of esophagus (p63+, KRT5+, KRT8−, KRT7−) and stomach (P63−, KRT5−, KRT8+, KRT7+) respectively (Figure 2G-H, Figure 1C-H, Supplementary Figure 1C).

**Figure 2.**
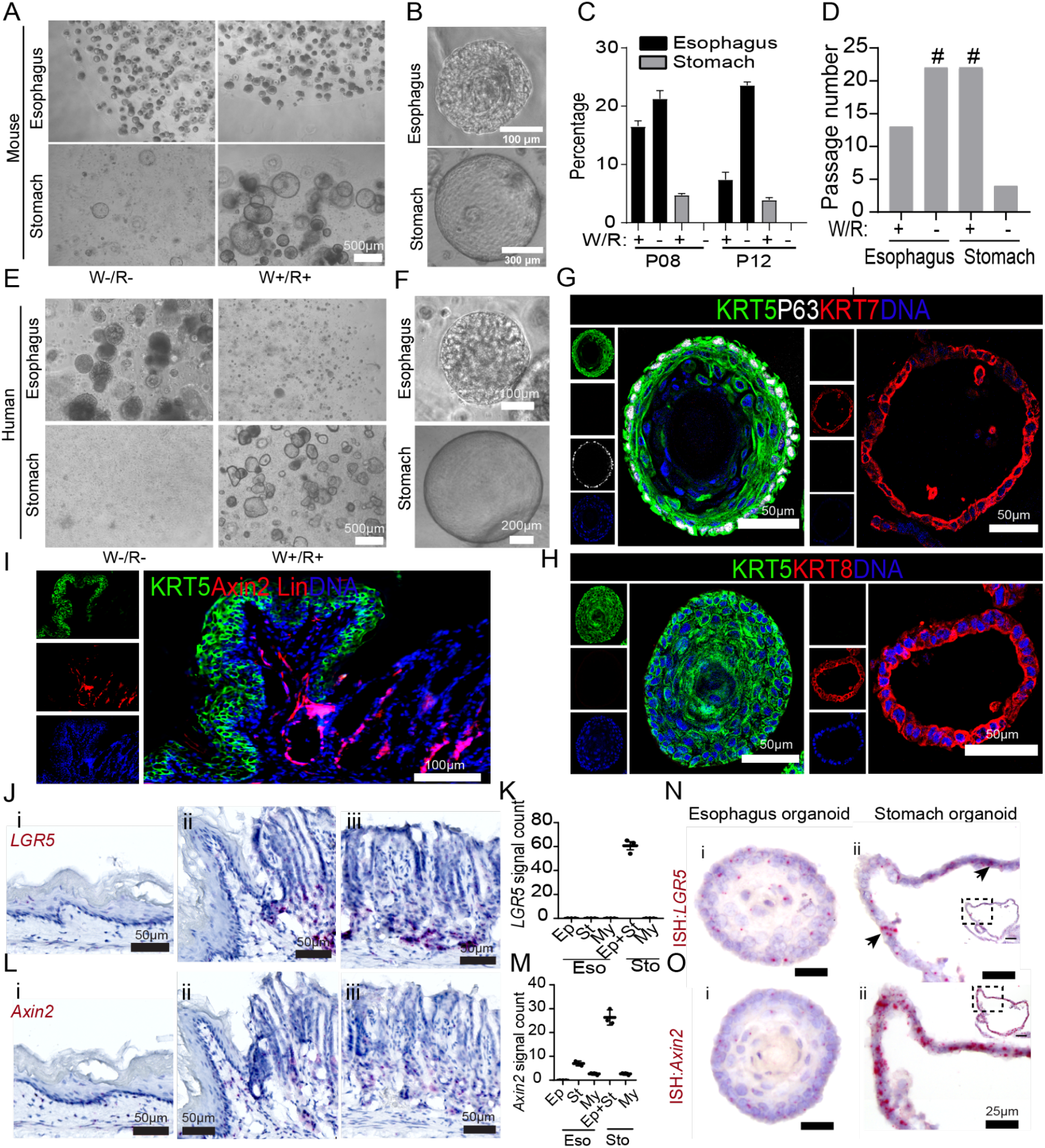
Esophageal stratified and columnar stomach organoid growth depend on the distinct Wnt microenvironmental factors. (A) Bright-field images of mouse squamous esophageal and columnar stomach organoids grown in the presence or absence of WNT3A (W) and R-spondin1 (R). (B) Higher magnification bright-field images of esophagus and stomach organoids grown in W−/R− and W+/R+ media, respectively. (C) Percentage of organoid formation from esophagus and stomach epithelial stem cells grown either in W−/R− and W+/R+ culture at indicated passage (P) number. n= mean+/−SD of three biological replicates. (D) Long term passaging of esophageal and stomach organoids in W−/R− and W+/R+ culture media. ‘#’ indicates organoids retained passaging ability beyond the indicated number. (E-F) Bright-field images of human squamous esophagus and columnar stomach organoids grown in W+/R+ or W−/R− media (E) and higher magnification bright-field images of the human esophagus and stomach organoid grown in W−/R− and W+/R+ media, respectively (F). (G-H) Confocal images of esophageal organoid (left panel) and stomach organoid (right panel) immunolabeled for KRT5 (green), KRT7 (Red), P63 (white) (G), KRT5 (green), KRT8 (Red) (H) and nuclei in blue. (I) Tiled images of GEJ sections from 17 weeks old *Axin2-Cre^ERT2^/Rosa26-tdTomato* mice after tamoxifen induction at the age of 4 weeks. Squamous epithelial cells were immunostained with KRT5 antibody (green), Axin2 lineage traced cells marked by Tdtomato (red), nuclei in blue. (J-M) smRNA-ISH images for the Wnt pathway genes *Lgr5* (J) and *Axin2* (L) in the mouse esophagus tissue (i), stomach gland at GEJ (ii), and in stomach glands (iii). Nuclei in blue. Quantification of *Axin2* (K) and *Lgr5* (M) signal counts in epithelia (Ep), stroma (St), and myofibroblast (My) in the esophageal and stomach tissue regions. Signal counts were performed in three non-overlapping 100 μm^2^ area of images. (N-O) smRNA-ISH images of mouse esophageal (i) and stomach (ii) organoids probed with *Lgr5* (N) and *Axin2* (O). Nuclei in blue. Inset image showing the whole organoid, A black arrow pointing to *Lgr5* expressing cells in the stomach organoid. Images in A-B and E-O are representative of n= 3 mice or human donors.

Further, a known marker of stomach stem cells located in the base of the gland, Leucine-rich repeat-containing G-protein coupled receptor 5 (LGR5) that binds to WNTs and WNT agonists R-spondins to activate the Wnt pathway ^29^, was found not expressed in the esophageal epithelial cells (Figure 2J-K). *Axin-2* gene, a downstream target of the canonical Wnt-beta-catenin signalling pathway, is expressed at the base of stomach glands but not in the esophageal epithelium (Figure 2L-M). Lineage tracing of Axin2 in mice further confirmed that esophageal epithelial cells were negative for Axin2 lineage while the Axin2+ cells labelled the columnar epithelium of the stomach gland (Figure 2I). Consistently, smRNA-ISH of organoids confirmed that, unlike stomach epithelium, esophageal epithelium does not express *Lgr5 or Axin2* genes (Figure 2N-O). Thus, contrasting Wnt signals regulate the differential proliferation of esophageal squamous stratified and stomach columnar epithelial lineages at the GEJ.

### Subcellular composition and transcriptional signatures of the gastric and esophageal epithelium

To identify the regulatory signatures of squamous and columnar epithelium of GEJ, we performed a global transcriptomic analysis of the esophageal and stomach organoids. Among 34393 unique genes, 8030 genes were differentially regulated between columnar and squamous epithelium (Figure 3A, Supplementary Table 1-2). Gene ontology terms associated with the differentially expressed genes between the esophagus and stomach organoids showed enrichment of distinct pathways specific to the epithelial types (Figure 3B, Supplementary Table 3). Pathways related to the epidermal cell development, keratinocyte differentiation, transcription and translation, regulation of cell-cell adhesion were highly enriched in the esophageal epithelial cells. In the stomach epithelial cells, metabolic and catabolic processes related to lipid, fatty acids and ion transport were enriched. Corroborating to the enriched GO terms, periodic acid–Schiff (PAS) staining of organoids and the GEJ tissue showed intense PAS staining in the columnar epithelium, indicating high expression and secretion of glycoproteins, glycolipids and mucins compared to the stratified epithelium (Figure 3C). Pathways related to epithelial cell proliferation, cell junction assembly, cell-substrate adhesion were similarly regulated between two epithelial lineages. Thus, these distinct organoids recapitulate the structural, functional and molecular similarity to the tissue of origin.

**Figure 3.**
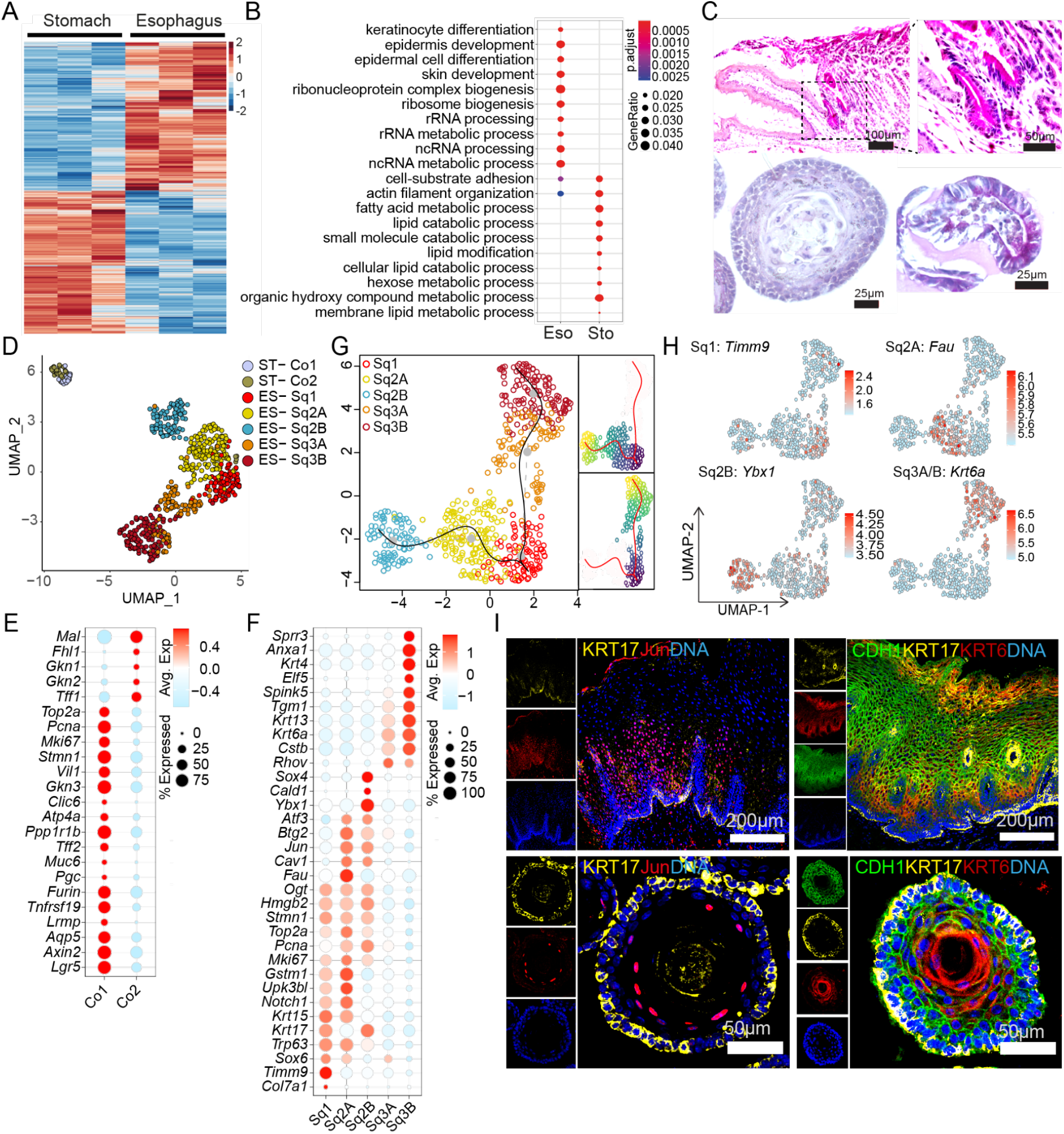
Bulk and single-cell transcriptomics of esophageal and stomach organoids reveal cellular subpopulation and lineage-specific signatures. (A) Heat map showing differentially expressed genes (DEG) in esophagus versus stomach organoids. Columns represent organoids derived from individual mice. The colour bar represents z-scored gene expression. (B) Top 10 enriched gene ontology (GO) terms associated with DEG between esophageal and stomach organoids. (C) PAS staining of mouse GEJ tissue section (i), esophageal organoid (ii) and stomach organoid (iii). (D) UMAP of scRNA-seq data derived from esophageal and stomach organoids showing cellular subclusters of each epithelium. Single cells are colour coded by cluster annotation (ST, stomach; ES, esophagus; Sq, squamous). (E-F) Dot plot depicting the expression of selected marker genes specific for stomach (E) and esophagus (F) epithelial subclusters. Circle size indicates the percentage of cells expressing indicated genes. Fill colour depicts the normalised and scaled mean expression levels from high (red) to low (blue). (G) UMAP showing the reconstruction of pseudo time trajectories in esophagus epithelial subclusters originating from Sq1. (H) Normalized expression values of selected markers colour coded on UMAP representing esophageal epithelial subclusters. (D-H) n= 3 biologically independent experiments. (I) Confocal images for the human tissue (upper panel) and mouse esophagus organoids (lower panel), stained with KRT17, Jun, KRT6 and CDH1. Nuclei were stained with DAPI (blue). Images represent 3 independent biological replicates.

Cytokeratins are intermediate filaments that enable cells to withstand mechanical stress and innate immunity and are uniquely expressed in different epithelial types ^32^. The analysis of the expression profile of the cytokeratins revealed that *Krt14, Krt15, Krt17, Krt5, Krt4, Krt13, Krt6a, Krt6b*, and *Krt16* were highly expressed in the squamous while *Krt8, Krt18, Krt7, Krt19, Krt20* are highly expressed in the columnar epithelium (Supplementary Figure 2A). While bulk transcriptomics using the esophagus and stomach organoids provided important insights into the epithelial-specific expression patterns and signalling networks, it does not reveal cell type-specific expression. Thus, we applied single-cell RNA sequencing (scRNA-seq) of the stomach and esophageal organoids to gain insights into cell type-specific gene expression patterns, cell states and the cellular developmental trajectories of the epithelium. The generated scRNA-seq data of the columnar stomach and stratified squamous esophageal epithelial cells were combined to perform unsupervised clustering by dimensionality reduction and visualisation by uniform manifold approximation and projection (UMAP). The UMAP plot separated cell populations into two major clusters, one containing the columnar stomach and the other containing the esophageal epithelial cells, revealing the distinct transcriptional profiles of these two epithelial types (Figure 3D, and Supplementary Table 4). Further, cluster analysis was performed to characterise the heterogeneity and identify subpopulations within each epithelial type. Based on this analysis, these two epithelial types were further divided into seven transcriptionally distinct subclusters. The columnar epithelial cells of stomach (ST) organoids were segregated into two distinct clusters (ST-Co1, ST-Co2). In comparison, the squamous epithelial cells of esophageal (ES) organoids were segregated into five unique clusters (ES-Sq1, ES-Sq2A, ES-Sq2B, ES-Sq3A and Sq3B) (Figure 3D, and Supplementary Table 4). We observed that UMAP was able to recapitulate the differentiation stages of the columnar stomach and stratified esophageal epithelial cells. The ST-Co1 subcluster was enriched for the expression of well-known stomach stem cells markers such as *Lgr5*, *Aqp5* and *Axin2 together with* high levels of *Pgc*, *Muc6*, *Gkn3* and *Atp4a* expression, which are key markers of cells present in the neck and isthmus region. These cells also expressed high levels of proliferation markers, including *Mki67, Pcna, Top2a* and *Stmn1*. The second subcluster of stomach cells, termed ST-Co2, were found to express high levels of, *Gkn1, Gkn2, Tff1* representing the pit cells of the gastric gland (Figure 3E and Supplementary Table 4) ^33^. In contrast to the stomach, where the markers for different types of cells within the gastric gland, including stem cells and different differentiated cells, are well characterised, the knowledge regarding the characteristics of esophageal stem cells and the feature of the differentiating cells is minimal. The ES-Sq1 subcluster expresses *Col7a1, Timm9, Trp63, Stmn1*, and *Krt17*, representing the stratified epithelium’s basal cells. The ES-Sq2A subcluster was enriched for expression of *Fau, Gstm1, Jun* in addition to *Upk3lb, and* several proliferation markers, including *Mki67, Top2a, Pcna*. The subcluster ES-Sq2B was enriched for *Atf3*, *Cav1, Ybx1, Cald1* and *Sox4*. The ES-Sq3A and ES-Sq3B subclusters exhibited a gradual increase in the expression of genes such as *Rhov*, *Krt6a, Krt13*, *Anxa1*, *Tgm1*, *Spink5*, *Gsta5, Sprr3* and *Elf5* (Figure 3F, 3H, Supplementary Figure 2B-E, and Supplementary Table 5). To further understand the spatial pattern of expression of these esophageal subcellular marker proteins, we verified their expression patterns by the Human Protein Atlas (HPA) database ^34^. The expression of proteins of the ES-Sq1 subcluster is mainly restricted to the basal cells of the esophagus epithelium. ES-Sq2A and ES-Sq2B expressed markers of the parabasal cells, excluding the basal and fully differentiated cells. The ES-Sq3A subcluster expressed markers predominantly expressed in differentiated cell layers above parabasal cells, while the ES-Sq3B subcluster genes marked the terminally differentiated layers of the esophageal epithelium. Together, our data indicate that the ES-Sq1 with KRT17^hi^/Jun^Low^ population constitute the stem cells of the healthy esophagus epithelium from which other subclusters arise by differentiation. To further validate gene expression dynamics and assign the progression of the stem cells and the path of differentiation of their descendants, we performed a pseudo-temporal reconstruction of the lineage structure using the slingshot lineage inference tool ^35^. Pseudotemporal modelling facilitates the reconstruction of differentiation trajectories based on gene expression transition when cells change from one state to the next. This analysis revealed two distinct trajectories, all originating from the basal stem cell compartment of ES-Sq1 (Figure 3G). This was further validated by immunostaining for the KRT17, Jun and KRT6 in human and mouse tissue and organoids, revealing exclusive KRT17+/JUN− basal stem cells while the parabasal cells above expressed both KRT17 and JUN and the differentiated cells expressed high levels of Krt6 (Figure 3H).

### Transcriptional signatures identify divergence of canonical and non-canonical Wnt pathways in gastroesophageal epithelia

While Wnt signalling was critical in regulating the GEJ homeostasis (Figure 1H-J, K-M, 2J-K, L-M), much less is known about the non-canonical, β-catenin-independent Wnt signalling in the gastroesophageal epithelium. Our analysis unravelled that columnar epithelial cells were enriched for the canonical Wnt beta-catenin pathway genes and non-canonical Wnt/Ca2+ pathway genes. In contrast, squamous epithelial cells were enriched for the non-canonical Wnt/planar cell polarity (PCP) pathway genes (Figure 4A). The genes coding for key proteins mediating canonical Wnt signalling *Axin2*, *Lrp5*, *Lrp6* and transcription factor *Tcf7* were highly upregulated in the columnar epithelium of the stomach. Also, the non-canonical Wnt/Ca2+ pathway genes, including *Camk2b, Camk2d*, and *Nfatc2* were upregulated in stomach organoids. In contrast, only non-canonical Wnt/PCP pathway genes *Scrib*, *Rac1*, *Serpinb5*, *Daam1* and *Vangl1* were highly expressed in the squamous epithelial cells of the esophagus (Figure 4A). We further identified the distinct expression patterns of the canonical and non-canonical Wnt signalling genes in the different subpopulations of the columnar and esophageal epithelium from the scRNA seq data (Figure 4B). We found that PCP pathway genes are predominantly expressed in the differentiated cells in the squamous epithelium of the esophagus. Thus, transcriptional signatures of the squamous and columnar organoids recapitulate the difference in the stem cell characteristics, tissue structure, and diverged function of epithelial tissue of GEJ.

**Figure 4.**
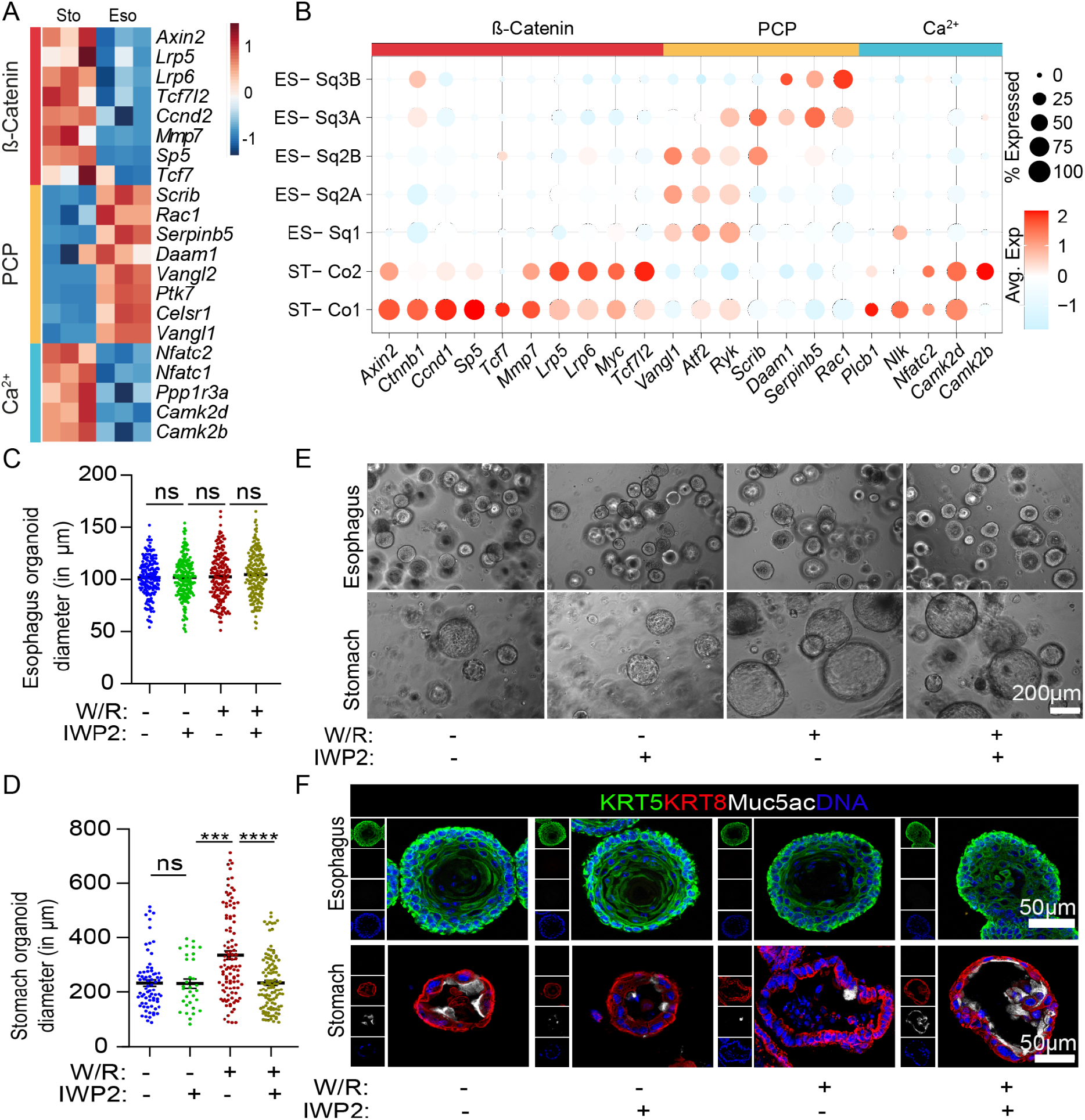
Canonical and non-canonical Wnt signaling in gastroesophageal epithelial regeneration. (A) Heat map showing expression of differentially regulated Wnt signalling pathway genes in esophagus versus stomach organoids. Columns represent organoids derived from 3 individual mice. The colour bar represents z-scored gene expression. (B) Dot plot depicting expression of canonical and non-canonical Wnt pathway associated genes in the stomach and esophagus epithelial subclusters. Circle size indicates the percentage of cells expressing an indicated gene. Fill colour depicts the normalised and scaled mean expression levels from high (red) to low (blue). (C-F) Mouse stomach and esophagus organoids were grown in W+/R+ or W−/R− culture medium, additionally either treated or untreated with 5 μM WNT secretion inhibitor IWP2. Organoid size in diameter was measured for esophageal squamous organoids, n ≥ 183 (C); and stomach columnar organoids, n ≥ 32 (D). n= number of organoids measured from 3 biological replicates. ns= Non significant, *** =p<.01, ****=p<.0001. (E) Bright-field images representing the esophagus and stomach organoids. (F) Confocal images of esophageal organoid and stomach organoid immunolabeled with KRT5 (green), KRT8 (Red), Muc5ac (white), and nuclei stained with DAPI (blue). Images in E and F are representative of n= 3 mice.

Further, we analysed the endogenous Wnt signalling in the regulation of stomach and esophageal epithelial stemness and differentiation. Stomach and esophagus organoids were grown in the presence and absence of W/R conditioned media. In addition, organoids were treated with pan canonical and non-canonical WNT secretion inhibitor IWP2 ^36^. Treatment of IWP2 did not influence growth and the size of KRT5+ stratified organoids grown in the presence or absence of W/R conditioned media (Figure 4C, E and F (upper panel)). However, the absence of W/R conditioned media and additional treatment with IWP2 led to growth inhibition of the KRT8+ columnar organoids (Figure 4D, E and F (lower panel). The addition of IWP2 to W/R conditioned media to stomach organoids led to smaller organoids than control, and most of the cells showed accelerated differentiation with high expression of Muc5Ac (Figure 4F). Our data demonstrate that KRT5+ stratified epithelial stem cell maintenance and regeneration are WNT independent while both canonical and non-canonical WNT signalling play critical roles for KRT8+ columnar epithelial stemness and differentiation.

### Wnt signalling and gastroesophageal epithelial plasticity

Next, we investigated if the squamous and columnar epithelial types of GEJ originate from common or distinct adult stem cells and if they possess the plasticity to transdifferentiate between stratified and columnar epithelium in the presence or absence of Wnt growth factors. We induced lineage tracing in *Krt5-Cre^ERT2^;Rosa26-tdTomato* and *Krt8-Cre^ERT2^;Rosa26-tdTomato* mice (Figure 5A). Cells marked for KRT5 traced for the squamous epithelium only at the GEJ and in the entire region of the esophagus but not adjacent columnar epithelial cells (Figure 5B). In contrast, KRT8 marked cells traced exclusively in the columnar epithelial cells of the stomach (Figure 5C). Thus, suggesting that two distinct epithelial stem cells give rise to KRT5+ squamous lineage and KRT8+ columnar lineage that meet at GEJ. Next, we asked if these distinct epithelial stem cell lineages possess plasticity to transdifferentiate with altering Wnt growth factors. For this, epithelial cells from the esophagus and stomach were isolated from induced *Krt5-Cre^ERT2^;Rosa26-tdTomato* and *Krt8-Cre^ERT2^;Rosa26-tdTomato* mice and cultured as organoids in the presence or absence of W/R conditioned media. Irrespective of the presence or absence of W/R esophageal stratified organoids from *Krt5-Cre^ERT2^;Rosa26-tdTomato* mice were found to be labelled, whereas matched stomach columnar organoids were not (Figure 5D). Similarly, stomach columnar organoids from *Krt8-Cre;Rosa26-tdTomato* mice were found to be labelled, whereas matched esophageal stratified organoids were not labelled (Figure 5E). Further, immunofluorescence analysis confirmed that either presence or absence of Wnt factor did not change the expression of squamous specific marker KRT5 and columnar marker KRT8 in both epithelial organoid types (Figure 4F). Thus, the adult GEJ consists of two committed squamous and columnar epithelial stem cells that do not transdifferentiate with the change in the WNT microenvironment.

**Figure 5:**
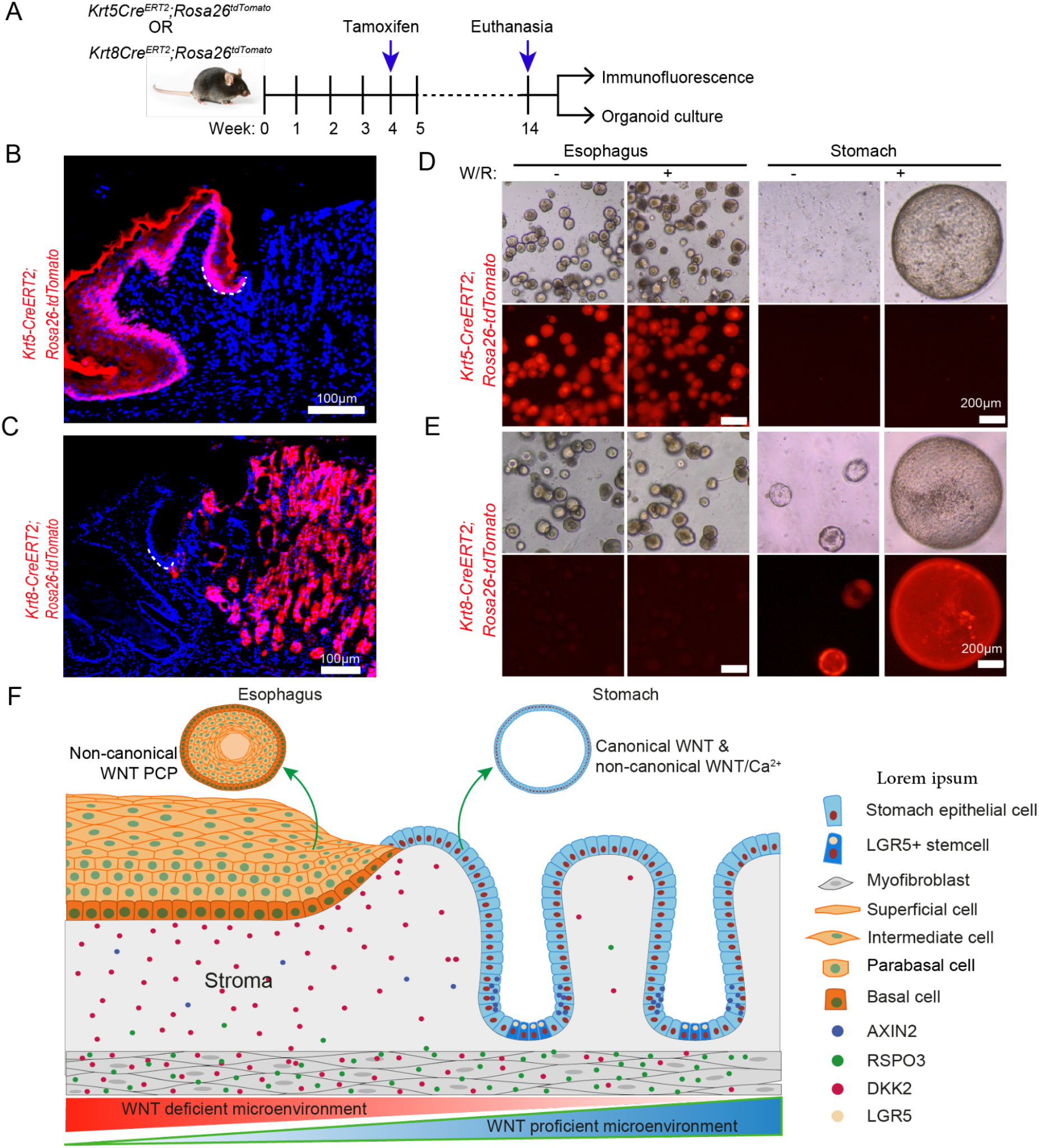
KRT5+ esophageal and KRT8+ stomach epithelial stem cells do not transdifferentiate under altered Wnt microenvironmental conditions. (A) Diagram representing the treatment scheme for lineage tracing of mice either expressing *Krt5-Cre^ERT2^;Rosa26-tdTomato* or *Krt8-Cre^ERT2^;Rosa26-tdTomato*. Cre recombinase was induced in mice by administering tamoxifen intraperitoneally at the age of 4 weeks on two consecutive days. Mice were euthanised in the 14th week, gastroesophageal tissues were either fixed for immunofluorescence or used to isolate esophagus and stomach epithelial cells to culture organoids. (B-C) Tiled images of GEJ from tissue sections of 14 weeks old *Krt5-Cre^ERT2^;Rosa26-tdTomato* (B) and *Krt8-Cre^ERT2^;Rosa26-tdTomato* (C) after Tamoxifen induction at the age of 4 weeks. Nuclei stained with DAPI (blue). The white dotted line indicates the basal cells of squamous epithelial cells in the esophagus near GEJ. (D-E) Organoids cultured in the absence and presence of (W/R) in the culture media from the lineage traced mice expressing either *Krt5-Cre^ERT2^;Rosa26-tdTomato* epithelial (D) and *Krt8-Cre^ERT2^;Rosa26-tdTomato* epithelial (E). (F) Schematic representation of distinct epithelial lineages and the underlying microenvironment in normal GEJ homeostasis. Data in B-E are representative of n= 3 mice.

## Discussion

Adult mucosal tissue regeneration and maintenance depends on the balanced action of tissue-specific stem cells self-renewal, proliferation, differentiation and cell-fate commitment. The tissue microenvironment, including stromal and immune cells, is critical in regulating stem cell regeneration and maintaining normal homeostasis ^2,33,37,38^. During tissue injury, the tissue microenvironment reprograms for the restoration of damaged tissue ^39^. However, if the damage-inducing stimuli persist, the tissue might develop adaptive phenomena such as metaplasia to cope with the stimuli ^40^. The gastroesophageal junction (GEJ) shows increased predisposition to the development of Barrett’s metaplasia (BE), enrichment of pathogenic microbes and carcinogenesis ^41,42^. This study unravelled the stage at which the GEJ border observed in normal adults is established during development, the epithelial cell types, plasticity and mechanisms regulating the normal GEJ homeostasis. These insights are invaluable for identifying the mechanism that deviates from normal tissue regeneration and homeostasis, contributing to disease onset.

Our systematic analysis of the epithelial lining of the developing esophagus and stomach from the embryonic stage to adult GEJ revealed that KRT8+/KRT7+ progenitors give rise to p63−/KRT8+/KRT7+ and p63+/KRT8+/KRT7+ cells, which eventually segregated by embryonic day 19 as distinct P63+/KRT5+/KRT8−/KRT7− squamous and P63−/KRT5−/KRT8+/KRT7+ columnar cell types at the GEJ and maintained in the adult. However, previous studies proposed that embryonic progenitors that uniquely express KRT7+ ^17^ or transitional p63+/KRT5+/KRT7+ ^18^ cells reside as few cells at the GEJ and are the cell of origin of BE. In contrast, we found that KRT7 RNA and protein are highly expressed in the stomach’s columnar epithelium and the basal cells of the esophageal squamous epithelium, albeit in lower levels but not restricted to a few GEJ cells. Our observation corroborates with other studies describing KRT7 expression in columnar gastric epithelial cells ^43^, while its absence does not alter the normal function of epithelial cells or influence other cytokeratin’s function ^44^. However, increased expression of KRT7 in BE ^43^ and other cancers ^45^ might be due to increased BMP4 signals, known to regulate KRT7 expression ^46,47^. Thus, we show the existence of two distinct epithelial lineage-specific stem cells that give rise to the squamous epithelium and columnar epithelium, which meet at the GEJ.

Signal crosstalk between the epithelium and underlying mesenchyme has been shown to direct the cellular differentiation and lineage specifications during embryogenesis ^48,49^. Here we found that in adult GEJ, spatially defined opposing Wnt signalling crosstalk establishes the borders at the squamocolumnar junction and their regeneration. Wnt signalling regulating morphogen, RSpondin 3 (Rspo3) from myofibroblasts of muscularis mucosa underlying the gastric glands in the antral region of the stomach are known to regulate adult gastric stem cell regeneration ^28^. Similarly, we found *Rspo3* expression in myofibroblasts underlying the glandular epithelium at the GEJ. Interestingly, unlike gastric glands where stem cells are proximal to *Rspo3* expressing muscularis mucosa, the basal stem cells of the esophagus and myofibroblasts are separated by wider lamina propria comprising stromal cells expressing higher levels of Wnt signalling inhibitor *Dkk2*. Consistently, growth factors inducing Wnt signalling were found to inhibit the development and long-term maintenance of stratified squamous organoids from the esophagus while supporting the development and stemness of both human and mouse gastric organoids. Thus, spatially restricted differential expression of Wnt signalling regulators underlying the adult GEJ epithelium establishes the borders. Corroborating to our observations, esophageal specification and separation from trachea during development is governed by the induction of WNT inhibitor molecules by the mesenchymal Barx1 ^50,51^. Likewise, similar principles were found to regulate uterine cervical squamocolumnar junction homeostasis ^2^.

Comparative analysis of bulk and single-cell transcriptional profiles of esophageal and gastric organoids revealed the subcellular composition and unique properties of these epithelial tissues with distinct cytokeratin profile and their divergence in function. Squamous stratified epithelium is associated with structural regulation, including keratinocyte proliferation and differentiation, cell-cell junctions, RNA biogenesis, while columnar epithelium is associated with metabolism and catabolism of fatty acids, lipids and polysaccharides. The results from scRNAseq suggest that squamous stratified and columnar organoids recapitulate the subcellular composition of the native esophageal and gastric epithelial tissue. This data further revealed the detailed molecular composition of different subcellular compartments of the esophagus epithelium, showing that the KRT17^hi^/Jun^low^ cells are the stem cells of the esophagus epithelium.

Besides developmentally associated canonical Wnt/beta-catenin pathways, Wnt signalling also comprises less characterised non-canonical Wnt/Ca2+ pathway and Wnt/PCP pathways. We found that columnar gastric epithelium shows active canonical Wnt/beta-catenin pathways and non-canonical Wnt/Ca2+ signalling, while the non-canonical Wnt/PCP pathway is predominantly active in the stratified squamous epithelium of the esophagus. Inhibition of both canonical and non-canonical pathways in gastric epithelium revealed that extrinsic canonical Wnt is essential for the proliferation of the stem cells. In contrast, endogenous non-canonical Wnt/Ca2+ is essential for maintaining stemness and preventing differentiation of stem cells into MU5Ac foveolar pit cells. We found that Wnt/PCP signalling implicated in tissue morphogenesis and epithelial cell polarity during embryogenesis is particularly active in the parabasal cells and not essential for squamous stratified stem cell regeneration and differentiation. Moreover, alterations in the Wnt signalling did not induce transdifferentiation between columnar or squamous type epithelia. Upregulated Wnt signalling is observed in BE compared to squamous lineage at the GEJ ^27^. However, the observed high Wnt signalling in BE could be due to differential outgrowth of columnar lineage in the esophagus.

In conclusion, we show that the adult stratified esophageal and columnar stomach epithelia and their subpopulations arise from distinct unilineage stem cells. The spatially defined antagonistic Wnt morphogen from the tissue microenvironment promotes differential proliferation of stomach and esophageal stem cells, thus maintaining the healthy GEJ homeostasis. Furthermore, our organoid models recapitulated the subcellular heterogeneity of the parent tissue and proved to be a powerful tool to model healthy tissue homoeostasis and disease development. Thus, these fundamental insights pave the way to understand the mechanisms underlying the development and progression of pathologies at the GEJ.

## Methods

### Mice

This study is compliant with all relevant ethical regulations regarding animal research. Animal research procedures were approved by the national legal, institutional and local authorities at the Max Planck Institute for Infection Biology. All animals were maintained in autoclaved micro isolator cages and provided sterile drinking water and chow ad libitum. Four- to twenty-week-old female mice were used for this study. Wild-type C57BL/6, *Krt5Cre^ERT2^* ^52^, and *Krt8Cre^ERT2^* ^53^ mice were obtained from the Jackson Laboratory. *Krt5Cre^ERT2^* and *Krt8Cre^ERT2^* strains were bred to *Rosa-tdTomato* mice ^54^ to generate mice expressing a fluorophore in Cre-expressing cells. For inducing Cre recombinase for the *Krt*5 or *Krt*8 lineage tracing, mice were administered with tamoxifen (Sigma) intraperitoneally (0.25 mg per g body weight in 50 μl corn oil) at week 4 for two consecutive days. Mice were euthanised at 14-20 weeks, and the gastroesophageal region was removed for further analysis. Experiments were performed in at least three biological replicates per condition. Mice were randomly allocated to experimental groups in all experiments.

The whole stomach was isolated from the mice at embryonic days 13, 16, 19, or postnatal mice were either used to isolate cells for organoid culture or fixed with 4% PFA for 1 hr at RT. Dehydration of the embryonic stomach was performed by immersing tissue with a series of ethanol, isopropanol, and acetone for 20 min each, followed by embedding with paraffin.

#### Antibodies and Chemicals

The following antibodies and chemicals were used: mouse anti-E-Cadherin (BD Biosciences, 610181), mouse-anti-E-Cadherin-488 (BD Biosciences, 560061), rabbit anti-cytokeratin 5-Alexa488 (Abcam, ab193894), mouse anti-p63 (Abcam, ab375), rabbit anti-cytokeratin 7 (Abcam, ab181598), rabbit anti-cytokeratin 7-Alexa555 (Abcam, ab209601), rabbit anti-cytokeratin 8 (Abcam, ab59400), mouse anti-MUC5AC (Abcam, ab212636), rabbit anti-cytokeratin 17 (Abcam, ab109725), mouse anti-c-Jun (Abcam, ab280089), mouse anti-cytokeratin 6 (Abcam, ab18586), donkey anti-rabbit–Cy3 (Jackson Immuno Research, 711-166-152), donkey anti-rabbit–Alexa Fluor 647 (Jackson Immuno Research, 647 711-605-152), donkey anti-mouse–Cy5 AffiniPure (Jackson Immuno Research, 715-175-151), Hoechst (Sigma, B2261), Draq5 (Cell Signaling, 4085), DAPI (Roche, 10236276001), IWP2 (Tocris Bioscience, 3533).

### Organoid culture and maintenance

#### Epithelial stem cell isolation from the human gastroesophageal junction

Human esophagus and stomach and Z line (GEJ) samples were provided by the Department of Hepatology and Gastroenterology, Charité University Medicine, Berlin, Germany. Usage for scientific research was approved by their ethics committee (EA4/034/14); informed consent was obtained from all subjects. The study is compliant with all relevant ethical regulations regarding research involving human participants. Tissue biopsies from anonymous donors were processed within 2–3h after removal. Biopsies were sourced from standard procedures.

#### Esophageal organoids

Human and mouse esophageal tissue was washed with sterile PBS and was cut open longitudinally, and minced with a sterile scissor into small pieces, transferred to a 15 ml centrifuge tube containing warm 3 ml 0.5 mg ml^−1^ collagenase type II (Calbiochem, 234155) solution and incubated for 30 min at 37°C shaker at 180 rpm. The tissue was mechanically disrupted with a 1 ml pipette tip by pipetting up and down ten times and centrifuged at 1000g. Pellet was resuspended with warm 3 ml TrypLE express (Gibco, 12604021), incubated for 30 minutes at 37°C in a shaker at 180 rpm. The tissue was mechanically disrupted with a 1 ml pipette tip by resuspending up and down ten times and passed through a 70 μm cell strainer (Falcon, 352350) to filter out larger tissue debris. Isolated cells were washed once with ADF++ media and resuspended with 50 μl of Matrigel (Corning, 356231) and plated on a pre-warmed 24 well plate. After polymerisation of Matrigel for 10 minutes at 37°C, matrigel was overlaid with complete 3D esophageal media containing ADF medium supplemented with 12 mM HEPES, 1% GlutaMax, 1% B27, 1% N2, 50 ng ml^−1^ murine EGF (Invitrogen, PMG8043), 100 ng ml^−1^ murine noggin (Peprotech; 250-38-100), 100 ng ml^−1^ human FGF-10 (Peprotech, 100-26-25), 1.25 mM N-acetyl-L-cysteine, 10 mM nicotinamide, 2 μM TGF-β R kinase Inhibitor IV, 10 μM ROCK inhibitor (Y-27632), 10 μM forskolin (Sigma, F6886) and 1% penicillin/streptomycin (Gibco, 15140-12).

As mentioned above, for human organoid culture, esophagus cells were isolated using collagenase type II and TrypLE treatment. Isolated cells were cultured in the 3D esophageal medium with similar composition as above, where EGF and noggin were replaced with human forms with 10 ng ml^−1^ human EGF (Invitrogen, PHG0311), 100 ng ml^−1^ human noggin (Peprotech; 120-10C-1000).

### Stomach Organoids

Mouse stomach tissue was incubated with 0.5 mM DTT/3 mM EDTA in PBS for 90 minutes at RT. Tissue was transferred to the ice-cold PBS and shaken vigorously to isolate stomach glands. 100 glands were mixed with 50 μl of Matrigel and plated on a pre-warmed 24 well plate. After polymerisation of Matrigel for 10 minutes, Matrigel was overlaid with complete 3D stomach media containing ADF medium supplemented with R-spondin1 conditioned medium (25%) and Wnt3A-conditioned medium (25%), 12 mM HEPES, 1% GlutaMax, 1% B27, 1% N2, 50 ng ml^−1^ murine EGF, 100 ng ml^−1^ murine noggin, 100 ng ml^−1^ human FGF-10, 1.25 mM N-acetyl-L-cysteine, 10 mM nicotinamide, 2 μM TGF-β R kinase Inhibitor IV, 10 μM ROCK inhibitor (Y-27632), 10 mM gastrin and 1% penicillin/streptomycin.

For human stomach organoid culture, stomach gland cells were isolated using collagenase type II treatment for 30 min, as mentioned above. Isolated cells were cultured in the Matrigel using a 3D stomach medium with similar composition as above, where EGF and noggin were replaced with human forms with 10 ng ml-1 human EGF, 100 ng ml^−1^human noggin (10 μM SB202190 (Sigma, S7067).

#### Organoid-forming efficiency and size analysis

Epithelial cells were counted, and a defined number was resuspended in 50 μl of Matrigel to generate organoids as described above. One week after plating, images were acquired from the whole well, and the number and diameter of formed organoids were determined using ImageJ to calculate the organoid-forming efficiency and measurement of size.

#### Immunofluorescence and microscopy

Mouse gastroesophageal tissue was cut longitudinally from the antrum through the greater curvature of the stomach to the esophagus. Tissue was fixed with 4% PFA overnight at RT, dehydrated by passing through a series of ethanol, isopropanol, xylene treatment for 60 minutes in a Leica TP1020 tissue processor and embedded with paraffin. Organoids were removed from Matrigel by washing five times with ice-cold PBS and fixed with 4% PFA for 1h at RT. After washing with PBS, dehydration of organoids was performed by a series of ethanol, isopropanol, and acetone treatment for 20 min each, followed by paraffinisation.

For immunofluorescence staining, 5 μm paraffin sections were cut on a Microm HM 315 microtome. The sections were deparaffinised, rehydrated, and treated with antigen retrieval solution (Dako, S1699). Sections were blocked using a blocking buffer (1% BSA and 2% FCS in PBS) for 1 hr at RT. Primary antibodies were diluted in blocking buffer, and sections were incubated overnight at 4°C, followed by five times PBS washes and 1 hr incubation with secondary antibodies diluted in blocking buffer along with Hoechst or Draq5. For direct fluorochrome tagged antibodies, sections were blocked with a blocking buffer for 1hr after adding a secondary antibody. Sections were washed with PBS five times and mounted using Mowiol. Images were acquired with a Leica TCS SP8 confocal microscope, or tiled images were obtained with an AxioScan.Z1 tissue imager (Zeiss), processed with Zen 2.3 (Blue edition) and compiled with Adobe illustrator. Confocal images were processed with Adobe Photoshop and analysed by using Image J software.

GEJ tissue was fixed with 2% PFA for 1 hr at 4°C in the dark for staining lineage traced mice. Tissue was washed with PBS and freshly frozen using dry ice-cooled isopentane and OCT compound (Tissue Tek, 4583). 5 μm tissue sections were cut using Cryomicrotome, washed with PBS. Tissue sections were used for either nuclei staining or immunofluorescence, as mentioned in the method.

##### Single-molecule RNA in situ hybridisation (smRNA-ISH)

For single-molecule RNA in situ labelling, paraffin-embedded 10 μm tissue sections were used with RNAscope 2.5 HD Red Reagent kit (Advanced Cell Diagnostics). Hybridisations were performed according to the manufacturer’s protocol. In each experiment, positive (PPIB) and negative (DapB) control probes were used according to the manufacturer’s guidelines. Tiled bright-field images were acquired with Axio Scan.Z1 tissue imager (Zeiss). Images were further processed with Zen 2.3 (Blue edition) image analysis software and further compiled using Adobe illustrator.

#### RNA isolation and quality control for microarray analysis

Microarrays were performed from mouse organoids cultured from esophageal and stomach epithelial stem cells (n = 3 biological replicates). Organoids were washed with ice-cold PBS and were pelleted and resuspended in 1 ml Trizol (Life Technologies), and RNA was isolated using a kit according to the manufacturer’s protocol. Quantity of RNA was measured using a NanoDrop 1000 UV-Vis spectrophotometer (Kisker), and quality was assessed by Agilent 2100 Bioanalyzer with an RNA Nano 6000 microfluidics kit (Agilent Technologies).

#### Microarray expression profiling

Microarray experiments were performed as single-colour hybridizations on Agilent-028005 SurePrint G3 Mouse GE 8×60K, and Agilent Feature Extraction software was used to obtain probe intensities. The extracted single-colour raw data files were background corrected, quantile normalised and further analysed for differential gene expression using R and the associated BioConductor package LIMMA ^55^. To compare esophagus and stomach gene expression, we performed an unpaired t-test. Genes with a p-Value < 0.05 and log2 fold change of − 0.5849625 and 0.5849625, corresponding to a 1.5-fold decrease, or increase in abundance, respectively, were considered differentially expressed. All statistical analysis was performed with R unless stated otherwise.

#### Overrepresentation analysis of microarray data

We performed an over-representation analysis (OA) on genes significantly higher expressed in the stomach or esophagus with gene sets based on GO biological process gene annotations ^56^. The analysis was performed in R using the function compareCluster from the package ClusterProfiler ^57^. As input, we took all significantly differentially expressed genes with a valid Entrez ID, which are 3234 genes higher expressed in the stomach and 3415 genes higher expressed in the esophagus. We used the default setting of ClusterProfiler as significance cutoff, an adjusted p-Value < 0.05 adjusted with the Benjamini-Hochberg-method [https://www.jstor.org/stable/2346101].

#### Single-cell preparation for scRNA-seq

Organoids were harvested with ice-cold PBS, pelleted by centrifugation (5 Min, 300g, 4°C), and Matrigel was removed. The process was repeated twice to remove Matrigel completely. Organoids were then incubated with warm TrypLE in a shaker (15 min, 37°C, 180 rpm). Organoids were sheared using a 1 mL pipette by pipetting up and down 20 times. Dissociated cells were passed through a 40 μm cells strainer to obtain suspension of single cells, and the cells were washed with 0.1% BSA in 1XPBS.

#### Multiplexing individual samples for scRNA-seq

Following the preparation of single-cell suspension, multiplexing of samples was performed according to the MULTI-seq protocol ^58^. The cells were counted, and a total of 1×10^6^ cells/sample were resuspended and pelleted at 1000g for 5min. The pellet was resuspended in 180μL of 3X SSC buffer with 1%BSA. To this 20μl of 20X lipid-modified DNA oligonucleotide (LMO) anchor: unique “sample barcode” oligonucleotides mix (20X= 4μM) in order to be multiplexed, with each sample receiving a different sample barcode. Samples were then incubated on ice for 5min. Then samples were supplemented with 20μl of 20X (20X= 4μM) common lipid-modified co-anchor to stabilise the membrane residence of barcodes. Samples were incubated on ice for an additional 5min. Barcode-containing media was then removed by adding 500μl of ice-cold 3X SSC containing 1% BSA to the samples and pelleted at 1000g for 5min at 4°C. The resulting cell pellet was washed again with 500μl of ice-cold 3X SSC+1% BSA, and the pellet was resuspended in 150μl of ice-cold 0.125X SSC + 0.04% BSA. The resuspended cells were counted, samples were pooled together equally, and cell numbers adjusted to 1000 cells/μl.

#### scRNA-seq library preparation and MULTIseq

A 10x Chromium Controller was used to partition single cells into nanolitre-scale Gel-Bead-In-EMulsions (GEMs). Approximately 2500 cells per sample were pooled and loaded onto the controller. Single-cell suspensions were processed using the 10x Genomics Single Cell 3′ v3.1 RNA-seq kit. Reverse transcription, cDNA amplification and construction of the gene expression libraries were prepared following the detailed protocol provided by 10x Genomics. After the cDNA Amplification step, the MULTIseq barcode fraction was separated and processes according to the MULTIseq protocol ^58^. A SimpliAmp Thermal Cycler (Applied Biosystems) was used for amplification and incubations. Libraries were quantified by Qubit™ 3.0 Fluorometer (ThermoFisher), and quality was checked using a 2100 Bioanalyzer with a High Sensitivity DNA kit (Agilent). Sequencing was performed in paired-end mode with an S1 100-cycles kit using Novaseq 6000 sequencer (Illumina).

#### Processing of raw sequencing data

Sequencing data was processed using the CellRanger (v3.1.0) pipeline from 10x Genomics. Generation of FASTQ files for both gene expression and MULTI-seq libraries was achieved from the raw sequencing data using the “cellranger mkfastq” command with default parameters. We then used “cellranger count” with default parameters to perform alignment against the mm10 build of the mouse genome, UMI counting and for generating the feature barcode matrix.

#### scRNA-Seq sample De-Multiplexing

In order to determine the sample origin of each cellular barcode, the generated MULTI-seq FASTQ files were processed using the R package deMULTIplex (v1.0.2)^58^. The resulting sample barcode UMI count matrix data was fed as the input for MULTI-seq sample classification, by which cells from the same sample were grouped. Suspected cells that were positive for more than one sample barcode were classified as doublets. In general, sample multiplexing is not a perfect process in which small groups of cells can remain ‘negative’ without falling into any of the sample groups.

Therefore, a semi-supervised negative cell reclassification was performed using the functions ‘findReclassCells’ and ‘rescueCells’ to rescue the negative cells. These rescued (re-classified) cells were added back to their respective predicted sample groups. Finally, each cell containing the information regarding the sample group (including negatives and doublets) was utilised for scRNA sequencing downstream analysis.

#### single-cell RNA-seq data quality control, normalisation and clustering

The obtained filtered gene expression matrix was analysed using R software (v.4.0.3) with the Seurat ^59^ package (v.4.0.0). The demultiplexed sample barcode UMI information was incorporated into the gene expression matrix. We chose to omit unrescued cells (negatives and doublets) from the data for further analysis, resulting in 1099 cells. As a next step, we scrutinised for potential doublets by neglecting barcodes with less than 100 genes, more than 8500 genes and more than 80,000 UMI counts. Low-quality cells with more than 20% of the UMIs derived from the mitochondrial genome were excluded. Ultimately, we split each unique sample into a separate Seurat object based on the MULTI-seq sample barcodes, which contained 765 cells from the esophagus and 90 cells from stomach samples designated for further downstream analyses. Normalisation and variance stabilisation of these objects was performed using a negative binomial regression model provided by the sctransform ^60^ package (v.0.3.2), which also identified the highly variable genes. In addition, the mitochondrial mapping percentage and cell cycle scores (calculated using CellCycleScoring command) were regressed out during data normalisation and scaling. Dimensionality reduction of the data was performed using the RunPCA function with default parameters. Clustering was done using the FindNeighbors and FindClusters functions on the top 30 principal components, which was then visualised by implementing a nonlinear dimensionality reduction with the RunUMAP function. We identified a set of cells with erroneously annotated sample barcodes, which might be due to the negative cell reclassification during the demultiplexing process. Hence, we carefully assessed for the presence of such other cells, (e.g. mixup cells/doublets with substantial and coherent expression profiles of a hybrid transcriptome based on columnar and squamous epithelial marker gene expression (*K*rt*8/18* and *Krt5/14/6a/13*, respectively) and excluded them from further analysis. As an outcome, UMAP was derived from analysing a total of 612 cells from esophagus and stomach samples combined (Figure 3D). To further unravel the subpopulations present within the data, we reclustered the esophageal and stomach cell clusters separately by repeating the aforementioned workflow for dimensionality reduction and clustering.

#### Cell-Type Annotation and differential gene expression identification

Cells were projected into 2D space after performing dimensionality reduction and were clustered together based on their transcriptional similarities. The resulting cell clusters were annotated based on specific canonical marker genes (Figure 3E-F). Additionally, to identify genes that would discriminate these clusters, we used the FindAllMarkers command with default Wilcoxon rank-sum test in Seurat to identify the differentially expressed genes between cell clusters/type with default parameters (Supplementary Tables 4-5).

#### Trajectory Inference/Pseudotime Analysis

Developmental trajectories in the data were modelled using the Slingshot ^35^ package (v.1.6.1). We identified the global lineage structure using the minimum spanning tree (MST) approach provided by the getLineages function. This resulted in different smoothened lineages, which were modelled by fitting simultaneous principal curves using the getCurves function and also the information regarding how the cells are ordered in each lineage based on pseudotime values.

##### Statistics and reproducibility

GraphPad Prism (v.8) was used for statistical calculations and the generation of plots. The data are displayed as mean±s.e.m. P<0.05 was considered to be statistically significant. Each experiment was repeated independently with similar results.

#### Human Protein Atlas analysis of esophageal subcluster marker genes

To find the spatial distribution of our selected subcluster specific markers at the protein level (Figure 3F), we scanned the HPA database (http://www.proteinatlas.org, v20.1), which provides information on the tissue and cell distribution of human proteins based on immunostaining.

## Acknowledgements

We thank M. Drabkina, K. Hoffman for technical assistance; I. Wagner for the microarrays; D. Son for help with sample preparations. N.K. is supported by Deutsche Forschungsgemeinschaft Deutscher Akademischer Austauschdienst (DAAD) and Deutsche Forschungsgemeinschaft Graduiertenkolleg DFG-GRK2157, P.G.P is supported by the DFG-GRK 2157. C.C. is supported by University Würzburg and DFG-GRK 2157. The funders had no influence on the study design or analysis of the data.

## Author contributions

C.C., R.K.G. conceived the study; C.C. R.K.G. and N.K. designed the experiments, performed and analysed the data; N.K. and C.C. prepared the single cells for scRNA seq; S.M.K contributed to IHC experiments; T.K, C.T and A.-E.S performed the multiplexed scRNA-seq and raw data pre-processing; P.G.P performed scRNA-seq bioinformatics analysis, and C.W. performed microarray bioinformatics analysis with the help of C.C. N.K and R.K.G; V.B. contributed Axioscan imaging; H.-J.M. contributed microarray studies; T.F.M provided the infrastructure and advice; C.J., M.B. and B.W. provided human samples; N.K. R.K.G and C.C. wrote the manuscript.

---

**Figure S1.**
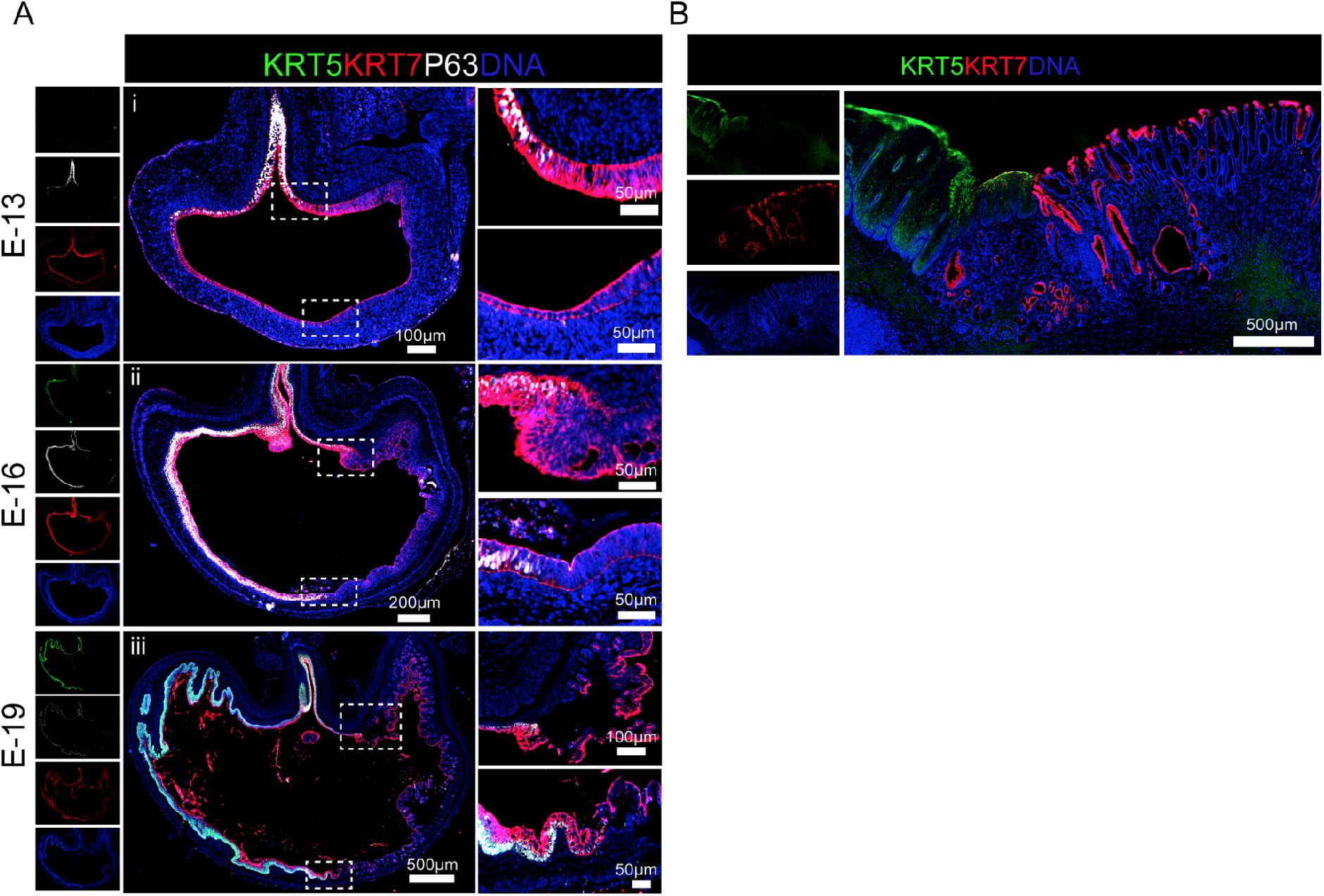
Distinct cytokeratins mark stratified squamous esophageal and columnar stomach epithelium. (A) Tiled images of tissue sections from mouse stomach from embryonic day 13 (E-13) (i), E-16 (ii), and E-19 (iii) stained with KRT5 (green), KRT7 (Red), P63 (white). Nuclei are stained with DAPI (blue). A magnified view of the boxed GEJ regions is shown in the right panel. (B) Tiled images of adult healthy human GEJ tissue sections stained with KRT5 (green), KRT7 (Red) and nuclei stained with DAPI (blue). Images are representative of n=3 mice or human donors. Tiled images were acquired with an AxioScan imager.

**Figure S2:**
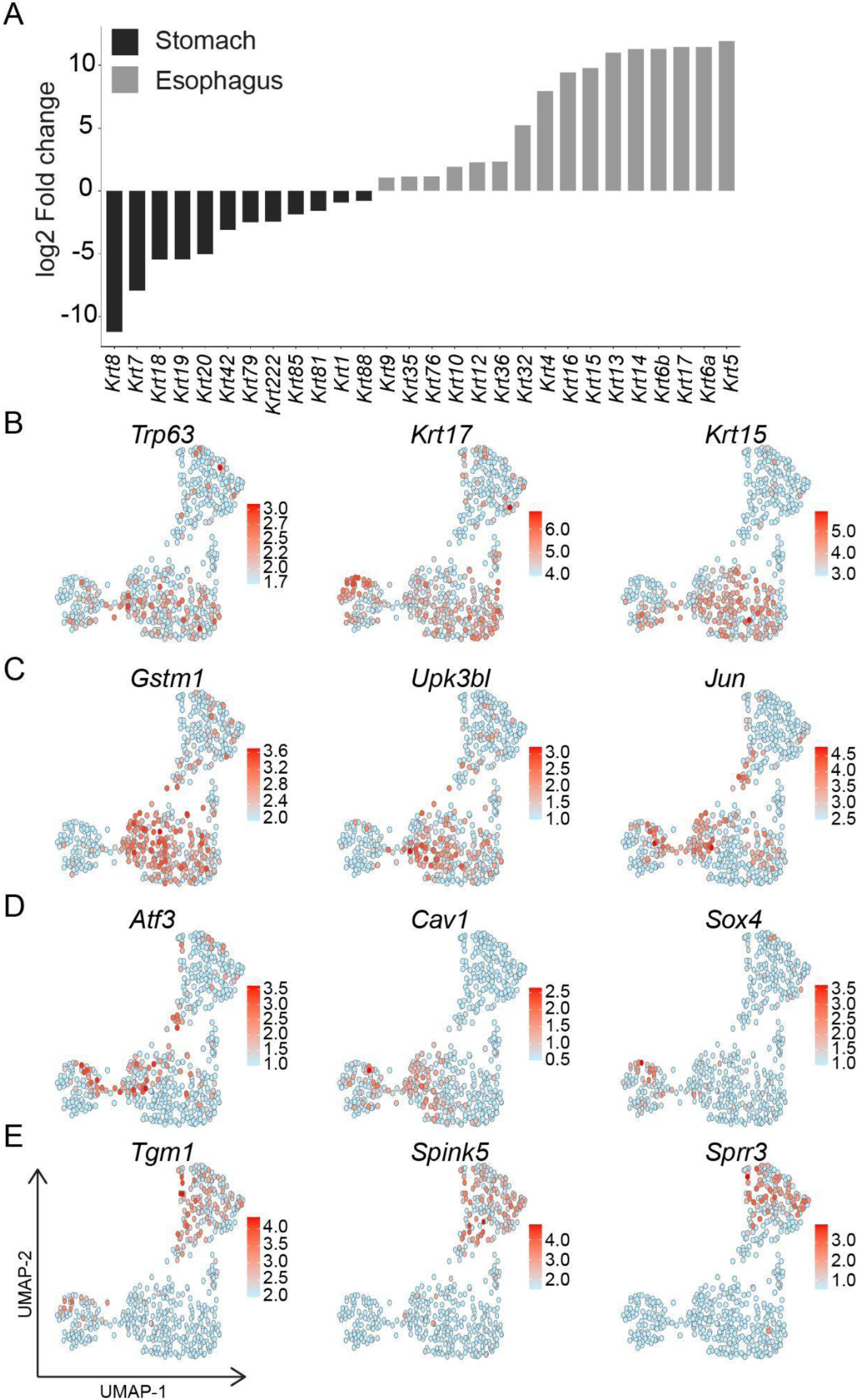
Distinct expression of genes in stomach versus esophagus and subclusters of esophagus epithelium. (A) Bar plot depicting Log2FC of differentially expressed cytokeratin genes between mouse esophageal and stomach organoids, revealing a distinct expression profile. (B-E) Normalised expression values of selected markers colour coded on UMAP representing esophageal epithelial subclusters as in Fig 3 D, F, G for Sq1 (B), Sq2A (C), Sq2B (D), Sq3A and Sq3B (E). n = 3 biologically independent experiments.

## Notes

### Competing Interest Statement

The authors have declared no competing interest.

